# Interpreting ciliopathy-associated missense variants of uncertain significance (VUS) in an animal model

**DOI:** 10.1101/2021.06.17.448799

**Authors:** Karen I. Lange, Sunayna Best, Sofia Tsiropoulou, Ian Berry, Colin A. Johnson, Oliver E. Blacque

## Abstract

**Purpose:** Better methods are required to interpret the pathogenicity of disease-associated variants of uncertain significance (VUS), which cannot be actioned clinically. In this study, we explore the use of a tractable animal model (*Caenorhabditis elegans*) for *in vivo* interpretation of missense VUS alleles of *TMEM67*, a cilia gene associated with ciliopathies.

**Methods:** CRISPR/Cas9 gene editing was used to generate homozygous knock-in *C. elegans* worm strains carrying *TMEM67* patient variants engineered into the orthologous gene (*mks-3*). Quantitative phenotypic assays of sensory cilia structure and function measured if the variants affect *mks-3* gene function. Results from worms were validated by a genetic complementation assay in a human *TMEM67* knock-out hTERT-RPE1 cell line that tests a *TMEM67* signaling function.

**Results:** Assays in *C. elegans* accurately distinguished between known benign (Asp359Glu, Thr360Ala) and known pathogenic (Glu361Ter, Gln376Pro) variants. Analysis of eight missense VUS generated evidence that three are benign (Cys173Arg, Thr176Ile, Gly979Arg) and five are pathogenic (Cys170Tyr, His782Arg, Gly786Glu, His790Arg, Ser961Tyr).

**Conclusion:** Efficient genome editing and quantitative functional assays in *C. elegans* make it a tractable *in vivo* animal model that allows rapid, cost-effective interpretation of ciliopathy-associated missense VUS alleles.

## INTRODUCTION

Exome and genome sequencing have revolutionised our ability to identify the genetic causes of disease. Missense variants (single codon altered to encode a different amino acid) are the most numerous class of protein-altering variants^1^ but only a subset are associated with disease^2^. Based on disease features and patterns of inheritance, identified variants are classified as benign, likely benign, uncertain significance, likely pathogenic or pathogenic (as defined by the Association for Clinical Genomic Science; ACGS)^3^. For novel or previously uncharacterized variants, the only evidence available to assess their pathogenicity is population allele frequency and analysis by *in silico* tools (e.g. SIFT/PolyPhen/CADD), which are not sufficient to meet the threshold for a ‘likely pathogenic’ classification according to best practices established by ACGS^3^. A VUS (variant of uncertain significance) classification is made when there is insufficient evidence to conclude on pathogenicity^3,4^. Currently, most missense variants are classified as VUS (116,215/198,080=58.7%, accessed from ClinVar^5^ September 2021). Since VUS classifications cannot be used for molecular diagnosis of disease, they cannot be acted upon clinically. Therefore, a VUS classification can delay or prohibit accurate disease management or genetic counselling, and prevents patients from accessing gene-specific therapies and clinical trials^6,7^. Given the wealth of genomic data available, there is a pressing clinical need to definitively reclassify VUS as benign or pathogenic. Effective experimental strategies for the functional interpretation of VUS are required to address this problem^8^. With the emergence of advanced genetics tools such as CRISPR-Cas9 gene editing, non-rodent model organisms such as zebrafish, *Drosophila* and *C. elegans* are emerging as robust *in vivo* experimental platforms for determining variant pathogenicity^9–11^.

Ciliopathies are a heterogenous group of at least 25 inherited disorders with clinically overlapping phenotypes, caused by pathogenic variants in almost 200 genes^12^. Ciliopathies are caused by defects in cilia, which are microtubule-based organelles, typically 2-20 microns long, that extend from the surfaces of most cell types. Motile cilia propel cells through a fluid or push fluid across a tissue surface. Primary cilia act as cellular “antennae”^13^, transducing a wide variety of extrinsic chemical and physical (eg. light, odorants) signals into the cell^14^.

Primary cilia are especially important for coordinating cell-cell communication signaling pathways (eg. Shh, Wnt, PDGF-a) essential for development and homeostasis^15^. Ciliopathies affect many organ systems, causing a broad range of clinical phenotypes of varied severity and penetrance that include cystic kidneys, retinal dystrophy, bone abnormalities, organ laterality defects, respiratory tract defects, infertility, obesity, neurodevelopmental defects and cognitive impairment^16^.

TMEM67 is a transmembrane protein that localizes specifically to the proximal 0.2-1.0 μm region of the ciliary axoneme called the transition zone^17,18^. Pathogenic variants in *TMEM67* cause several ciliopathies^19^ including Meckel Syndrome (OMIM #607361), COACH Syndrome (OMIM #216360), and are a major contributor to Joubert Syndrome^7,20^ (OMIM #610688). Most reported missense variants in *TMEM67* have uncertain or conflicting interpretations of their clinical significance (88/141=62.4%, accessed from ClinVar^5^ September 2021). The abundance of VUS alleles makes *TMEM67* an excellent candidate to explore if modelling VUS in *C. elegans* can generate evidence about their pathogenicity. We used CRISPR-Cas9 technology to model eight *TMEM67* missense VUS in *C. elegans*. Using quantifiable assays of cilium structure and function, we determined whether the variants are damaging or benign. We then validated the worm findings using a genetic complementation-based approach in *TMEM67* null human cells. Our study indicates that *C. elegans* is a tractable model system that can provide evidence of variant pathogenicity.

## METHODS

### Modelling of protein secondary structure

Human TMEM67 (NP_714915.3) and *C. elegans* MKS-3 (NP_495591.2) protein sequences were analysed by the RaptorX protein structure prediction server, using default settings for the deep dilated convolutional residual neural networks method^21^. Absolute model quality was assessed by ranking GDT (Global Distance Test) scores defined as 1*N(1)+0.75*N(2)+0.5*N(4)+0.25*N(8), where N(x) is the number of residues with estimated modelling error (in Å) smaller than x, divided by protein length and multiplied by 100. GDT scores >50 indicate a good quality model. However, the highest ranking models for TMEM67 (GDT = 28.968) and MKS-3 (GDT = 19.269) suggest that portions of these models are lower quality. Models in the .pdb format were visualized and annotated in EzMol (http://www.sbg.bio.ic.ac.uk/ezmol/).

### *C. elegans* maintenance

All *C. elegans* strains in this study were maintained at 20°C or 15°C on nematode growth medium seeded with OP50 *E. coli* using standard techniques ^22^. All worm strains are listed in Table S2.

### CRISPR/Cas9 to engineer *mks-3* mutants in *C. elegans*

CRISPR protocols were performed as previously described^23^ in *nphp-4(tm925)* using an *unc-58* co-CRISPR strategy. crRNA are listed in Table S3. The CRISPR efficiency (defined as the percent of F1 pools that were positive for the edit by PCR) varied from 1-35% with an average 15%. Accuracy of the engineered variants was confirmed with Sanger sequencing. *unc-58* was also sequenced and unintended *unc-58* mutations^24^ were outcrossed. PCR and sequencing primers are listed in Table S4.

### *C. elegans* quantitative phenotyping assays

Dye filling, roaming, and chemotaxis assays were performed as previously described^25^ with young adult hermaphrodites at 20°C. Assays were performed blinded to genotype with at least three independent biological replicates.

### Generating transgenic worms expressing extrachromosomal MKS-3::GFP

*mks-3::gfp* transgenes were generated with PCR-based fusion of *mks-3* gDNA (including 485 bp of 5’ UTR sequence) with GFP (pPD95_77, gift from Andrew Fire, Addgene plasmid #1495). All primers are listed in Table S4. *mks-3(tm2547)* hermaphrodites were injected with 0.25ng/μl *mks-3::gfp* and 100ng/μl coel::dsRed (gift from Piali Sengupta, Addgene plasmid #8938) to generate extrachromosomal arrays (1-7 lines each). PCR confirmed the presence of *mks-3::gfp* in the stable extrachromosomal arrays.

### *C. elegans* wide-field imaging and quantification of fluorescence

Young adult hermaphrodites were immobilised on 4% agarose pads in 40 mM tetramisole (Sigma). Images were acquired with a 100x (1.40 NA) oil objective on an upright Leica DM5000B epifluorescence microscope and captured with an Andor iXon+ camera. Image analysis was performed with FIJI/ImageJ (NIH). MKS-3::GFP fluorescence was quantified as previously described^23^.

### Transmission electron microscopy

Young adult hermaphrodites were processed as previously described^25^. Briefly, worms were fixed in 2.5% glutaraldehyde (Merck) in Sørensen’s phosphate buffer (0.1 M, pH 7.4) for 48 hours at 4°C, post-fixed in 1% osmium tetroxide (EMS) for 1 hour, and dehydrated through an increasing ethanol gradient. Samples were treated with propylene oxide (Sigma) and embedded in EPON resin (Agar Scientific) for 24 hours at 60°C. Serial, ultra-thin (90 nm) sections of the worm nose tissue were cut using a Leica EM UC6 Ultramicrotome, collected on copper grids (EMS), stained with 2% uranyl acetate (Agar Scientific) for 20 min followed by 3% lead citrate (LabTech) for 5 min, and imaged on a Tecnai 12 (FEI software) with an acceleration voltage of 120 kV.

### Cell culture

Human hTERT-immortalized retinal pigmentary epithelial (hTERT-RPE1, American Type Culture Collection; ATCC) wild-type and crispant cell-lines were grown in Dulbecco’s minimum essential medium (DMEM)/Ham’s F12 medium supplemented with GlutaMAX (Gibco #10565018) and 10% fetal bovine serum (FBS). For selected experiments involving cilia, cells following passage were serum-starved in DMEM/F-12 media containing 0.2% FBS. Cells were cultured in an incubator at 37°C with 5% CO2.

### TMEM67 cloning, plasmid constructs and transfections

Full-length *H. sapiens* TMEM67 isoform 1 (RefSeq JF432845, plasmid ID HsCD00505975, DNASU Plasmid Repository) was cloned into pENTR223. The ORF was Gateway cloned into a C-terminal GFP-tagged Gateway pcDNA-DEST47 vector (ThermoFisher Scientific), sequence verified, and sub-cloned into pcDNA3.1 myc/HisA vector with HiFi cloning (New England Biolabs). We also introduced an endogenous Kozak sequence prior to the start site, the first 30 nucleotides of the main transcript that were missing from the DNASU sequence, and a GS linker between the ORF and myc tag. To generate TMEM67 variants, we used the QuikChange II XL Site-Directed Mutagenesis Kit (Agilent) according to the manufacturer’s protocol. Primer sequences are listed in Table S5. The final constructs were verified by sequencing. Cells at 80% confluency were transfected with plasmids using Lipofectamine 2000 (Life Technologies Ltd.) as described previously^26^.

### CRISPR/Cas9 genome editing in cell culture

GFP-expressing pSpCas9(BB)-2A-GFP (PX458) (gift from Feng Zhang, Addgene plasmid #48138). Three crRNAs targeting human TMEM67 (RefSeq NM_153704.5) were designed using Benchling (https://benchling.com), selected for the highest ranking on- and off-target effects. crRNAs were ordered as HPLC-purified oligos from Integrated DNA Technologies in addition to Alt-R CRISPR-Cas9 tracrRNA-ATTO550 conjugates. crRNA sequences are listed in Table S6. Lyophilised pellets were resuspended in Tris-EDTA (TE) buffer (Qiagen) to give 100mM stocks. crRNA and tracrRNA were mixed (1:1), and incubated in nuclease-free duplex buffer (30 mM HEPES, pH 7.5, 100 mM potassium acetate) to make 300nM guide RNA master mixes. Before transfecting into cells the crRNA:tracrRNA duplexes were incubated with Lipofectamine 2000 at an RNA:Lipofetamine ratio of 2:1 in 200μl/well Opti-MEM for 20 minutes. 1 ml of media from 6-well plate wells was removed, the transfection reagents applied, and the cells incubated overnight. Media was changed to fresh DMEM/F-12 with 10% FBS after 16 hours, and cells incubated for 48 hours. Following transfection, fluorescence-activated cell sorting (FACS) was performed to enrich cells expressing GFP and to produce clonal populations. 96 well plates (Corning) were treated with 200 μl of 4% bovine serum albumin (BSA) per well for one hour. Wells were then filled with 100 μl of filter-sterilised collection buffer (20% FCS, 1% penicillin-streptomycin, 50% conditioned media, 29% fresh DMEM/F-12 media). Transfected cells were prepared for FACS by removing media, washing in PBS, and treating with trypsin for 5 min before resuspending in filter-sterilised sorting buffer (1x Ca^2+^/Mg^2+^-free PBS, 5 mM EDTA, 25 mM HEPES pH 7.0). A 70 μm filter was used to disperse cells into 4% BSA-treated polystyrene FACS tubes. A BD Influx 6-way cell sorter (BD Biosciences) was used to index sort GFP-positive cells, calibrated against un-transfected control cells. When an abundance of GFP positive cells were present, the top 5% were targeted for index sorting. After sorting, cells were incubated for 3 weeks at 37°C with 5% CO2, with weekly checks for growing colonies.

### PCR and sequence validation of crispant cell-lines

To extract DNA from colonies within the 96-well plates, cells were washed with 1x Ca^2+^/Mg^2+^-free PBS and resuspended in 50 μl of DirectPCR Lysis Reagent (Viagen Biotech) containing 0.4 mg/ml Proteinase K (Sigma, # P4850). Suspensions were incubated at 55°C for 5 hours, followed by 85°C for 45 minutes. 1 μl of DNA extracts were used in PCR reactions. Primers are listed in Table S7. variants were identified by Sanger sequencing (GeneWiz Inc.) followed by analysis using the Synthego ICE v2^27^. The following clones were chosen for further study: clone 40, heterozygous for c.369delC (p.Glu124Lysfs*12); and clone 16 carrying biallelic variants [c.519delT]+[ c.519dupT] ([p.Cys173Trpfs*20]+[ p.Glu174*]). All variants were predicted to result in nonsense mediated decay by the Ensembl Variant Effect Predictor^28^. Clone 21 was a negative control cell-line that was mock-transfected, underwent FACS, but was verified to carry wild-type *TMEM67*.

### Whole cell extract preparation and western immunoblotting

Whole cell extracts containing total soluble proteins were prepared from hTERT-RPE1 cells that were transiently transfected with 1.0 μg plasmid constructs in 90 mm tissue culture dishes, or scaled down as appropriate. 10 μg total soluble protein was analysed by SDS-PAGE (4-12% polyacrylamide gradient) and western blotting according to standard protocols. Primary antibodies used: mouse anti-β actin (1:10000, clone AC-15, Abcam Ltd., Cambridge, UK); rabbit polyclonal anti-TMEM67 (1:500, 13975-1-AP; ProteinTech Inc., Rosemont, IL, USA); goat anti-ROR2 (1:1000, AF2064; R&D Systems Inc., Minneapolis, MN, USA). Appropriate HRP-conjugated secondary antibodies (Dako UK Ltd.) were used (final dilutions of 1:10000-25000) for detection by the enhanced chemiluminescence “Femto West” western blotting detection system (Thermo Fisher Scientific Inc., Rockford, IL, USA) and visualized using a ChemiDoc MP imaging system (BioRad Inc., Hercules, CA, USA). Ratios of active phosphorylated ROR2 : unphosphorylated ROR2 isoforms were calculated by quantitating band intensity using ImageLab 5.2.1 software (BioRad Inc.) for three biological replicates, as described previously^26^.

### Statistical analyses

All *C. elegans* statistical analyses were performed in Microsoft Excel with the Real Statistics Resource Pack Version 7.2 (www.real-statistics.com). A Shapiro-Wilk test determined if data were normally distributed. Statistical significance was determined with an ANOVA followed by Tukey’s *post hoc* (chemotaxis and GFP quantification) or Kruskal-Wallis followed by Dunn’s (roaming) or Schaich-Hammerle (dye filling) *post hoc* tests. For cell culture results, a normal distribution of data was confirmed using the Kolmogorov-Smirnov test (GraphPad Prism). Pairwise comparisons were analysed with Student’s two-tailed t-test using InStat (GraphPad Software Inc.). *, **, and *** refer to p-values of <0.05, <0.01, and <0.001, respectively.

## RESULTS

### Selection of *TMEM67* variants for analysis

TMEM67 variants were selected using two criteria: (i) conservation of the mutated amino acid in the worm orthologue (MKS-3) (**Figure S1**), and (ii) presence of an adjacent Cas9 PAM site in the *C. elegans* genome to facilitate CRISPR gene editing. Using these criteria, we modelled eight missense *TMEM67* VUS in *C. elegans mks-3* (**Figure 1A, Table S1**). We also included two known benign and two pathogenic variants as controls.

**Figure 1.**
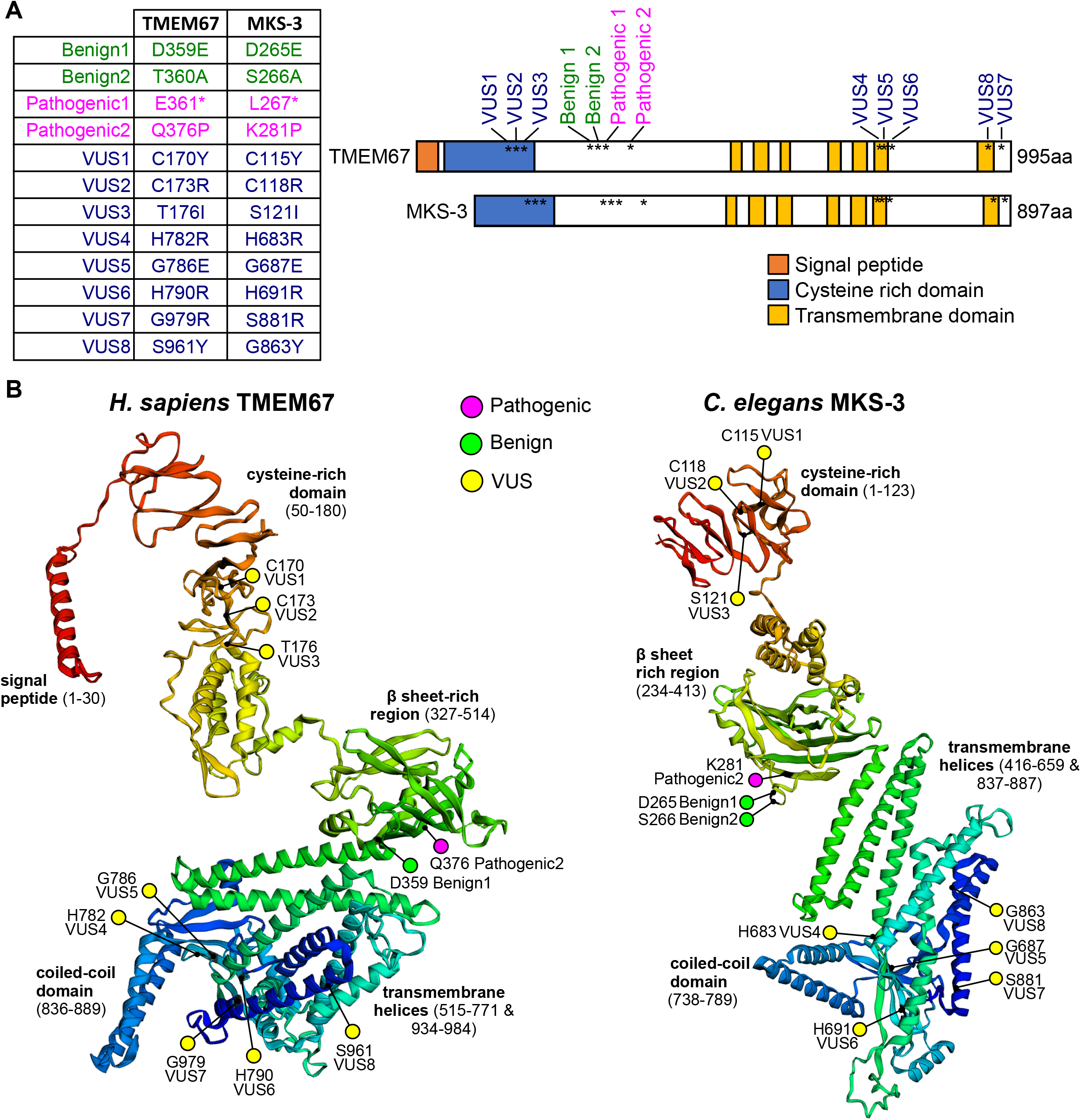
*TMEM67* variants analyzed in this study. A) Twelve variants in *TMEM67/mks-3* were generated by CRISPR-Cas9 gene editing and characterized in *C. elegans*. The schematic shows the relative positions of the variants along the length of the proteins. The domains are conserved between humans and worms, but the worm protein lacks the N-terminal signal peptide. B) RaptorX protein structure and domain organization predictions for the full-length TMEM67 and MKS-3 proteins. RaptorX is a deep learning algorithm that predicts secondary and tertiary structures of proteins without close homologues or known structures in the protein data bank. Ribbon diagrams of proteins are rainbow-coloured (red at N-terminus to dark blue at C-terminus) with variants indicated (magenta - known pathogenic; yellow - VUS; green - known benign) within the predicted protein domains. Note that transmembrane helices 1-5 (green to cyan) are followed by a single coiled-coil domain (light blue) and then two C-terminal transmembrane helices 6-7 (dark blue).

Benign1(Asp359Glu) and VUS4(His782Arg) were identified on the Ensembl variation database^29^. Benign2(Thr360Ala), Pathogenic1(Glu361Ter), Pathogenic2(Gln376Pro), VUS2(Cys173Arg), and VUS5(Gly786Glu) were identified on ClinVar^5^. VUS1(Cys170Tyr), VUS3(Thr176Ile), VUS6(His790Arg), VUS7(Gly979Arg), and VUS8(Ser961Tyr) were identified from clinical exome sequencing of Meckel syndrome foetuses. For simplicity, we refer to the variants using a shorthand notation (e.g. Benign1, Pathogenic1, VUS1). A *de novo* protein structure prediction program, Raptor X, revealed that the human and worm TMEM67 proteins show remarkable similarity in their overall predicted domain organization and secondary structure (**Figure 1B**). The targeted pathogenic, benign and VUS residues are present in comparable regions of secondary structure (**Figure 1B**). For example, the known benign variant residues are in exposed loops and the known pathogenic residues are buried in β-sheets (**Figure 1B**, **Figure S2**).

### Quantitative phenotypic analysis of *mks-3* VUS alleles in *C. elegans*

We employed a CRISPR/Cas9 genome editing strategy to engineer homozygous *mks-3* mutant worms (**Figure 1A**). Since *mks-3* functions redundantly with NPHP module genes (*nphp-1/4*), we generated the *mks-3* knock-in variants in an *nphp-4(tm925)* mutant background to facilitate phenotypic analysis^18^. *nphp-4(tm925)* is a 1109 bp deletion, subsequently referred to as *nphp-4(*Δ). In phenotypic assays, double mutant *mks-3(variant)*; *nphp-4(*Δ) phenotypes were compared to *mks-3(+)*; *nphp-4(*Δ) (wild-type, positive control) and *mks-3(*Δ); *nphp-4(*Δ) (949 bp deletion of *mks-3*, negative control)^18^. We hypothesized that pathogenic *mks-3* patient alleles would be phenotypically similar to the *mks-3(*Δ) allele.

To assess cilia structure and function we performed three quantitative assays: dye filling, roaming/foraging, and chemotaxis. The dye filling assay indirectly assesses the structural integrity of cilia^25^. We assessed lipophilic dye (DiI/DiO) staining in the four ciliated phasmid sensory neurons in the tail. Wild type and *mks-3(+)*; *nphp-4(*Δ) positive controls display robust dye filling, whereas the *mks-3(*Δ); *nphp-4(*Δ) negative control is dye filling defective (**Figure 2A**). Benign1 and Benign2-containing strains show robust dye uptake whilst strains with Pathogenic1 and Pathogenic2 are defective (**Figure 2A**). Five of the VUS alleles cause a severe dye filling defect (VUS1/4/5/6/8), whereas three (VUS2/3/7) do not (**Figure 2A**). *C. elegans* foraging behaviour is dependent on sensory cilia^25^. A single young adult worm is placed on a lawn of bacteria for 20 hours and the extent of its roaming across the plate is quantified (**Figure 2B**). *mks-3(+)*; *nphp-4(*Δ) positive control worms show a slight decrease in roaming compared to wild-type worms, whereas *mks-3(*Δ); *nphp-4(*Δ) negative controls exhibit a severe roaming defect (**Figure 2B**). As expected, Benign1 and Benign2-containing strains exhibit normal roaming behaviour, while Pathogenic1 and Pathogenic2-containing worm strains are roaming defective (**Figure 2B**). VUS4/5/6/8 exhibit a roaming defect, whereas worms containing VUS1/2/3/7 are roaming normal (**Figure 2B**). *C. elegans* chemotaxis towards benzaldehyde is also dependent on sensory cilia^25^. A population of 50-300 worms is placed in the center of a plate, equidistant from spots of control (ethanol) and benzaldehyde (1:200 in ethanol) solutions. *mks-3(+)*; *nphp-4(*Δ) positive control worms show a slight reduction in chemotaxis compared to wild type, whereas *mks-3(*Δ); *nphp-4(*Δ) negative controls exhibit a chemotaxis defective phenotype (**Figure 2C**). As expected, Benign1 and Benign2-containing strains are chemotaxis normal while Pathogenic1 and Pathogenic2-containing strains are defective (**Figure 2C**). VUS1/4/5/6-containing strains are severely chemotaxis defective, VUS3/8-containing worms exhibit an intermediate phenotype, and VUS2/7-containing worms are chemotaxis normal (**Figure 2C**).

**Figure 2.**
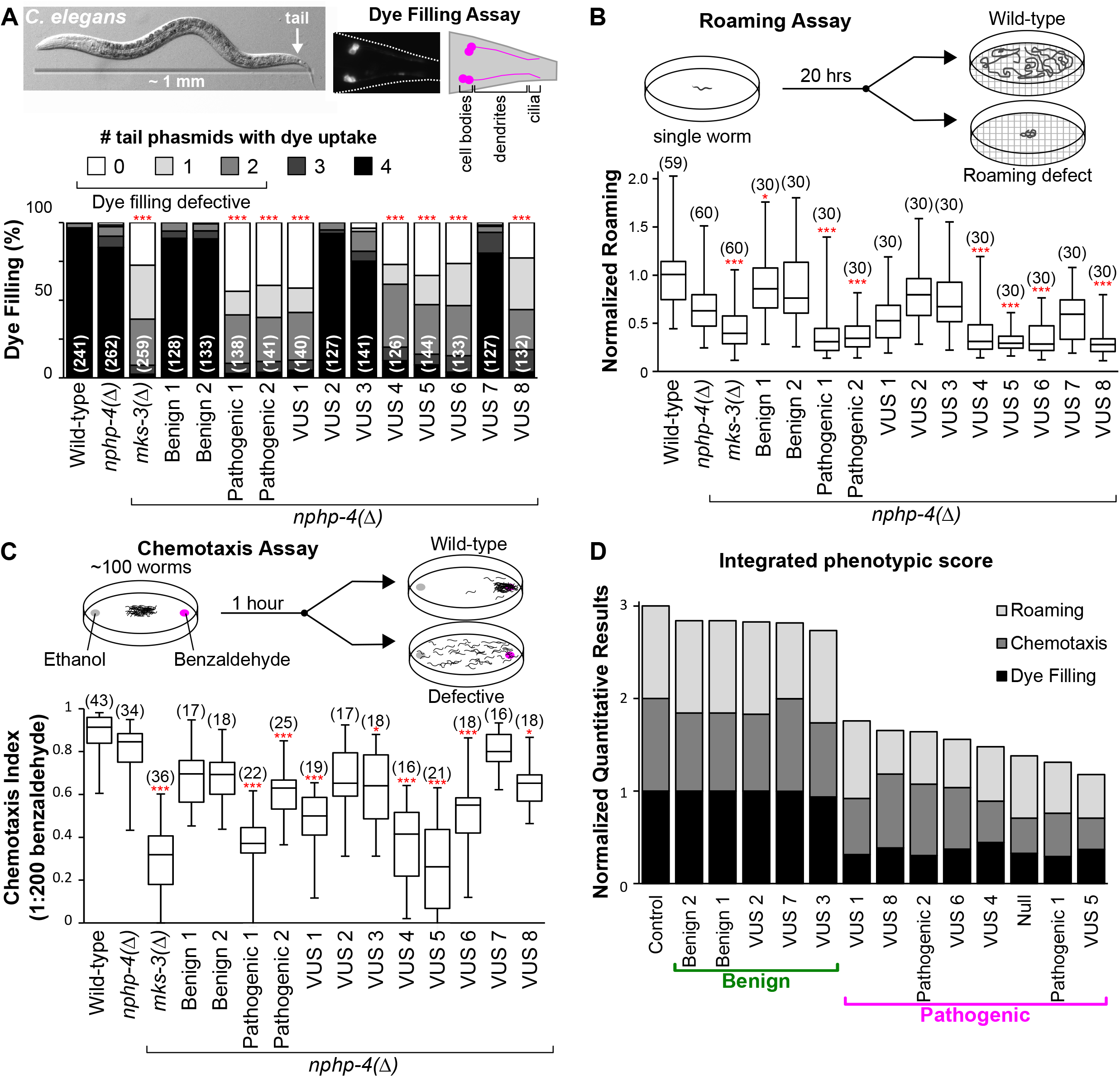
Quantitative phenotyping of cilia-dependent phenotypes in *C. elegans*. All assays were performed blind with at least three independent biological replicates. A) Lipophilic dye filling assay of the four phasmid (tail) neurons. The number of cell bodies which uptake dye was counted (values range from 0-4). The bar graph indicates the proportion of the population with dye uptake in 0 to 4 phasmid neurons. The number of worms is shown in brackets. Statistical significance according to Kruskal-Wallis followed by Schaich-Hammerle *post hoc* test. B) Assessment of worm roaming behaviour normalised to wild-type. A single young adult hermaphrodite was placed on a food-rich plate for 20 hours and the roaming activity was quantified. The number of worms is shown in brackets. Box plots indicate the maximum and minimum values (bars), median, lower quartile, and upper quartile. Statistical significance according to Kruskal-Wallis followed by Dunn’s *post hoc* test. C) Quantification of worm chemotaxis towards benzaldehyde after 60 minutes. Assay is performed on a population of 50-300 worms. The number of assays is shown in brackets.Statistical significance according to ANOVA followed by Tukey’s *post hoc* test. Box plots indicate the maximum and minimum values (bars), median, lower quartile, and upper quartile. D) Integration of the phenotypic results from the three quantitative assays into one value. Averages from each assay were normalized to the *nphp-4* control (with a maximum score of 1.0 for each assay) and summed. Values were ranked from highest (benign) to lowest (pathogenic).

To derive a predictive ‘interpretation’ score for the VUS alleles, we integrated the results from the three phenotypic assays into a single value (equal weighting; averages normalized to the *mks-3(+)*; *nphp-4(*Δ) positive control; maximum score of 1.0 per assay) (**Figure 2D**). The Benign1 and Benign2 variants score similarly to the *mks-3(+)*; *nphp-4(*Δ) positive control, whereas Pathogenic1 and Pathogenic2 variants score similarly to the *mks-3(*Δ); *nphp-4(*Δ) negative control (**Figure 2D**). Strains with VUS2, VUS3, or VUS7 received high scores comparable to the benign variants, whereas those with VUS1, VUS4, VUS5, VUS6 or VUS8 received scores comparable to the pathogenic variants. Therefore, in *C. elegans*, we conclude that VUS2/3/7 are benign variants and VUS1/4/5/6/8 are pathogenic variants.

### Effect of VUSs on transition zone localization of MKS-3 and ultrastructure

To provide further insight into the damaging or benign nature of the *TMEM67* VUS alleles, we examined the effect of the variants on the subcellular localisation of MKS-3. In *C. elegans* sensory neurons, transmembrane MKS-3 localizes to the ciliary transition zone (TZ), which corresponds to the most proximal 1 μm of the ciliary axoneme^18^. Fusion PCR was used to generate linear *mks-3::gfp* fragments (**Figure S3A**) containing variants of interest. Constructs were then expressed as extrachromosomal arrays in *C. elegans*, and a fluorescent lipophilic red dye (DiI) employed to co-stain the ciliary membrane. Pathogenic1 was excluded from this analysis because it is a nonsense allele with a premature stop codon. As expected, MKS-3(+)::GFP, Benign1::GFP, and Benign2::GFP exhibit TZ localization (**Figure 3A, S3B**). Pathogenic2::GFP showed no detectable fluorescence at the TZ, consistent with the finding that the human Pathogenic2 variant (Q376P) disrupts TMEM67 plasma membrane localization in cell culture^30^. Our conclusion that VUS2, VUS3, and VUS7 are benign (**Figure 2D**) predicts that the proteins should localize normally. Consistent with this hypothesis,VUS2::GFP, VUS3::GFP, and VUS7::GFP display TZ localizations in transgenic worms (**Figure 3A, S3B**). In contrast, transgenically expressed VUS1::GFP, VUS5::GFP, VUS6::GFP, and VUS8::GFP show no significant TZ localisation, consistent with these variants being pathogenic. Interestingly, despite a predicted pathogenic classification (**Figure 2D**), VUS4::GFP was TZ-localized in most transgenic worms, although signal levels were reduced (**Figure 3A, S3B**).

**Figure 3.**
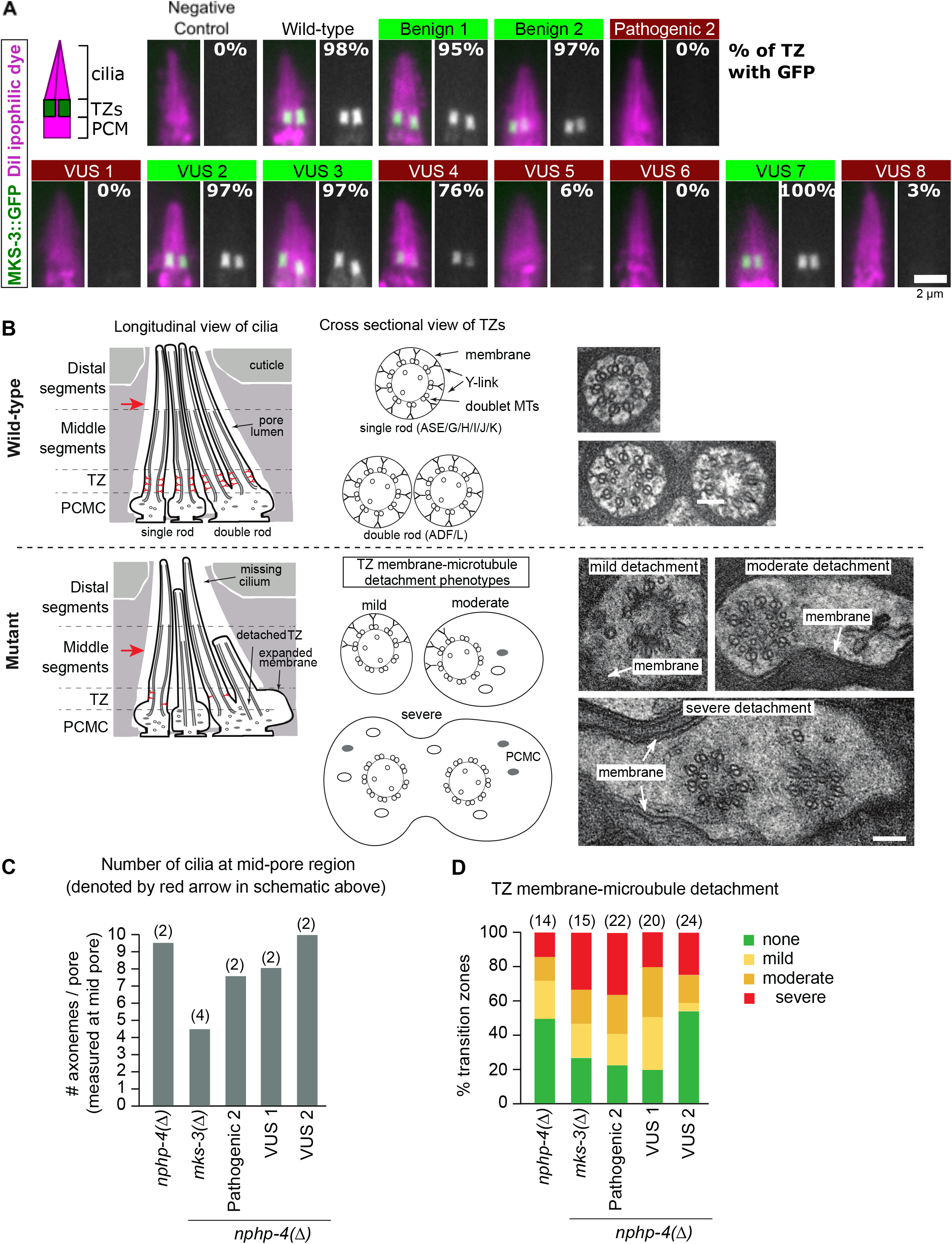
Testing *mks-3* variant predictions via analysis of GFP reporter localization and cilia/transition zone ultrastructure. A) Transgenic worm strains containing MKS-3::GFP extrachromosomal arrays were generated in *mks-3(*Δ) genetic background. DiI (magenta) lipophilic dye stains the cilia and peri-ciliary membrane (PCM). Grayscale image shows the green channel alone. The percentage of cilia with MKS-3::GFP localized to the transition zone (TZ) is indicated. Scale bar is 2μm. B) Ultrastructure of amphid channel ciliary axonemes and the TZ compartment in the indicated genotypes. Schematics show channel cilia in longitudinal orientation (only 4 of the 10 axonemes are shown for simplicity), and TZs in radial orientation. Electron micrograph images show the TZs in cross section. Scale bar is 100 nm. C) The mean number of cilia observed at the mid-pore region. The number of pores is shown in brackets. D) TZ membrane-microtubule detachment phenotypes of differing severity: mild - TZ microtubules are slightly detached from the membrane, which is expanded to a small degree; moderate - many TZ microtubules are clearly detached from the membrane, which is extensively expanded on one side of the axoneme; severe - most or all of the TZ microtubules are fully detached from the membrane, which is highly expanded, indicating that the TZ is ectopically docked in the periciliary membrane compartment (PCMC) cytoplasm. The number of TZs is shown in brackets.

We also assessed the effect of the TMEM67 VUS variants on the ultrastructure of cilia and TZs using transmission electron microscopy (TEM). Specifically, we analyzed two alleles: VUS1 (predicted pathogenic) and VUS2 (predicted benign). Pathogenic2 was also examined as a control. Wild-type amphidal pores contain 10 rod-shaped ciliary axonemes emanating from 8 sensory neurons (2 neurons possess a pair of rods), each consisting of middle (doublet microtubules) and distal (singlet microtubules) segments, and a proximal TZ compartment that emerges from a swelling at the distal dendrite tip called the periciliary membrane compartment (PCMC) (**Figure 3B**). At the middle pore region, *mks-3(+)*; *nphp-4(*Δ) positive controls display an almost full complement of 10 cilia, whereas at least two axonemes are missing in the *mks-3(*Δ); *nphp-4(*Δ) negative control and in Pathogenic2-containing worms (**Figure 3C**). We found that VUS1-containing worms are also missing some axonemes, whereas the VUS2-containing strain is not (**Figure 3C**). In cross-sections of the TZ and PCMC regions, approximately 50% of the TZs of *mks-3(+)*; *nphp-4(*Δ) positive controls show a wild type phenotype (TZ membrane and microtubules in close apposition, connected by electron dense Y-linkers) (**Figure 3B)**. As expected, the *mks-3(*Δ); *nphp-4(*Δ) negative control and the Pathogenic2-containing strain show a much more severe phenotype than *nphp-4(*Δ) alone (**Figure 3D**). The VUS2-containing strain displays a similar TZ phenotype to the positive control, whereas the VUS1-containing strain shows enhanced TZ membrane-MT detachment that is similar to the *mks-3(*Δ); *nphp-4(*Δ) negative control and Pathogenic2 (**Figure 3D**). Altogether, the ultrastructure data confirms the pathogenic and benign nature of VUS1 and VUS2, respectively.

### *In vitro* genetic complementation assay of TMEM67 VUS function in human cell culture

To further validate our findings from *C. elegans*, we utilized an *in vitro* human cell culture-based assay of *TMEM67* function. Previously, we demonstrated that TMEM67 is required for phosphorylation of the ROR2 co-receptor and subsequent activation of non-canonical Wnt signalling^26^ (**Figure 4A**). Here, we developed a genetic complementation approach using a validated hTERT-RPE1 crispant cell-line that has biallelic null mutations in *TMEM67* (**Figure S4**). In the absence of TMEM67, phosphorylation of ROR2 was not stimulated by exogenous treatment with the non-canonical ligand Wnt5a. Transient transfection with full-length wild-type TMEM67 fully rescued ROR2 phosphorylation following Wnt5a treatment. These responses allowed us to develop a series of assays to determine the relative effects of VUS on biological function. In this assay, transfection of Benign1 allowed 204.7% induction of phospho-ROR2 levels by Wnt5a relative to control (**Figure 4B**). In contrast, Pathogenic2 did not rescue biological function (90.2% induction) (**Figure 4B**). Comparison of all VUS, normalized to wild-type TMEM67 responses across three independent biological replicates, enabled us to interpret VUS1, VUS4, VUS5, VUS6 and VUS8 as pathogenic, and VUS3 and VUS7 as benign (**Figure 4C**). VUS2 was not tested in this assay.

**Figure 4.**
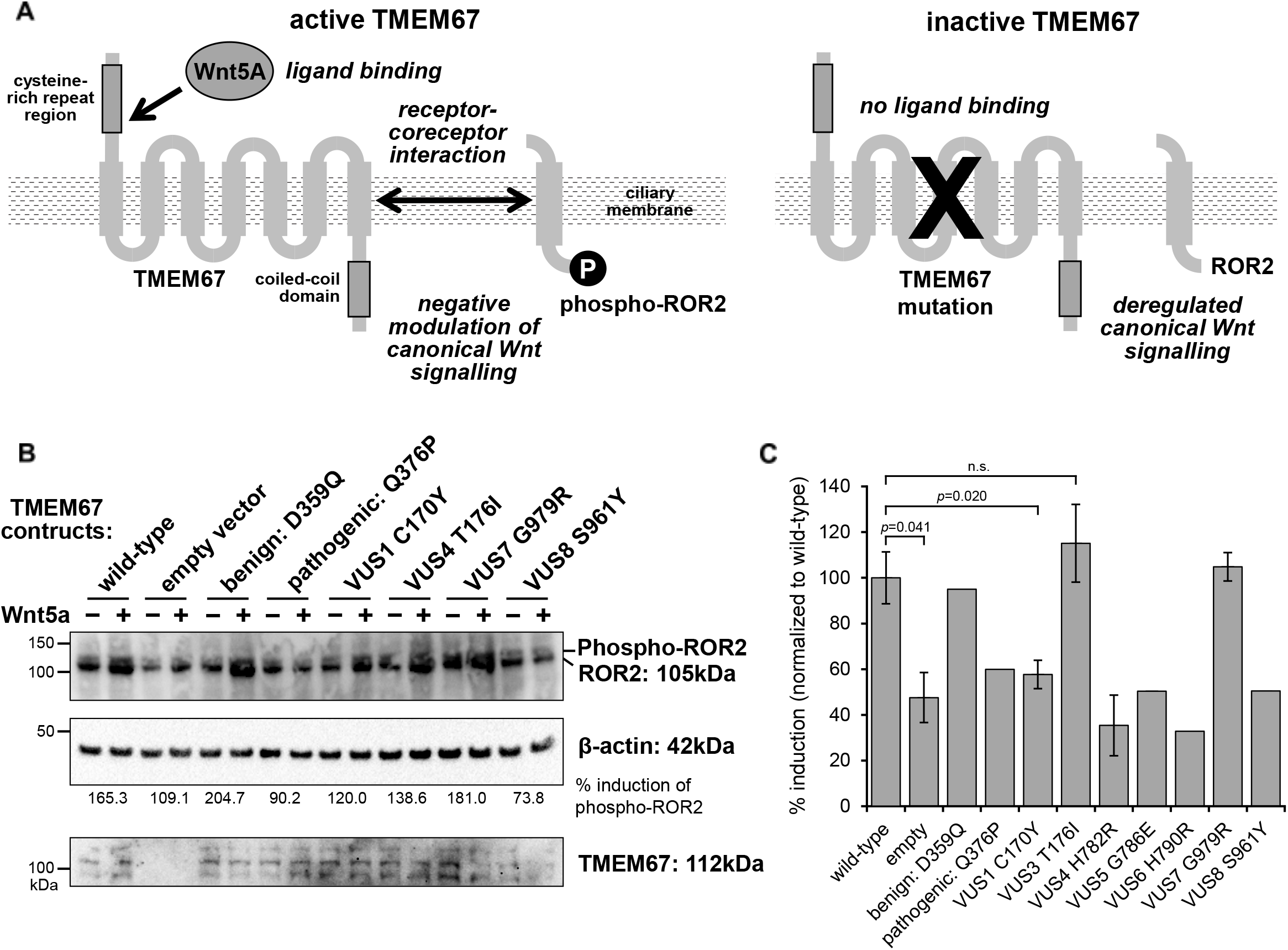
Validation of *C. elegans* predictions of TMEM67 VUS pathogenicity in cell culture. A) Schematic that summarizes the genetic complementation in hTERT-RPE1 *TMEM67* knock-out cells. Left: in the presence of TMEM67, phosphorylation of the co-receptor ROR2 is stimulated by exogenous treatment with the non-canonical ligand Wnt5a in comparison to control treatment. Right: if TMEM67 is lost or disrupted, ROR2 phosphorylation is not stimulated by this treatment. B) Western blots of ROR2, with upper phosphorylated isoform indicated (top panel), following transfection and expression (bottom panel) of TMEM67 constructs (wild-type, empty vector negative control, known benign variant control, known pathogenic variant control, and a selection of VUS alleles). Transfected *TMEM67^-/-^* knock-out cells were treated with control conditioned medium (-) or Wnt5a-conditioned medium (+). Loading control for normalization is β-actin. C) Densitometry scans of the phosphorylated ROR2 isoform (for n=3 biological replicates) were quantitated in the bar graph for % induction of Wnt5a-stimulated response compared to control response, normalized against responses to the wild-type TMEM67 construct. Pair-wise comparisons indicate statistical tests (Student t-test) for a minimum of n=3 biological replicates (n.s. not significant) with error bars indicating s.e.m. All other error bars indicate range values for n=2 replicates.

## DISCUSSION

Ciliopathies are multisystem disorders that affect many organs including kidneys, liver, and retina. While the organ systems affected by cilia dysfunction are not present in *C. elegans*, the basic biology of primary cilia is conserved. Despite undoubted context-specific distinctions, cilia proteins are functionally conserved^31,32^, therefore allowing us to model missense variants of these proteins in worms.

In this study, we exploited efficient genome editing and quantitative phenotypic analysis in *C. elegans* to determine the pathogenicity of *TMEM67* variants. This approach accurately classified known pathogenic and known benign variants. We also generated a pathogenicity prediction for all eight missense VUS alleles analyzed. Three VUS were phenotypically benign (VUS2(Cys173Arg), VUS3(Thr176Ile),VUS7(Gly979Arg)) and five were phenotypically pathogenic (VUS1(Cys170Tyr), VUS4(His782Arg), VUS5(Gly786Glu), VUS6(His790Arg), VUS8(Ser961Tyr)). These pathogenic predictions were validated in a human cell culture assay of TMEM67 function. We propose that experiments in *C. elegans* can interpret the pathogenicity of VUS and provide evidence towards their reclassification as benign or pathogenic.

Several *in silico* algorithms have been developed to predict the pathogenicity of missense variants. However, their accuracy is inconsistent^33–35^. Indeed, we tested five commonly used *in silico* tools (MISTIC, SIFT, Poly-Phen, CADD, and REVEL) and found that they return deleterious/damaging predictions for all 8 of the VUS examined in this project (**Figure S5**). The only exceptions are VUS1/3/8, where 1 or 2 of the tools returned non-deleterious or intermediate scores (**Figure S5**). Therefore, the algorithm predictions do not align with our observations in *C. elegans* and cell culture experiments. We conclude that *in vivo* modelling of missense variants in *C. elegans* more accurately predicts results in human cells than currently available prediction algorithms.

The quantitative assays employed in this study are suitable for high throughput analysis. Live animal fluorescence-activated cell sorting can be used for high throughput dye filling assays^36^ and automated worm tracking can be used for high throughput roaming and chemotaxis assays^37^. Additionally, machine learning can streamline the analysis of complex datasets to predict VUS pathogenicity^38^. One limitation to modelling patient variants in endogenous *C. elegans* genes is the conservation of amino acid residues. Although many cilia genes have clear orthologs in *C. elegans*, the amino acid conservation varies from 30-60%. *C. elegans* genes can be “humanized” to remove this limitation^38–40^. In summary, this study highlights that *C. elegans* is a practical model for variant interpretation of ciliary genes. Analysis of ciliopathy-associated VUS in *C. elegans* is accurate, quick, affordable, and easily interpretable. While this study focused on TMEM67, we anticipate that VUS alleles of any conserved ciliary genes can be modelled and characterized in *C. elegans*.

The authors declare no competing interests.

## Acknowledgements

This work was funded by Science Foundation Ireland (SFI) in partnership with the Biotechnology and Biological Sciences Research Council (BBSRC) under grant number 16/BBSRC/3394 (to OEB) and BB/P007791/1 (to CAJ). SB acknowledges support from a Wellcome Trust clinical training fellowship (203914/Z/16/Z). We thank Joseph McNicholl for assistance with construction of the *mks-3::gfp* transgenes.

