## Supplemental Data for "Interpreting ciliopathy-associated missense variants of uncertain significance (VUS) in an animal model"

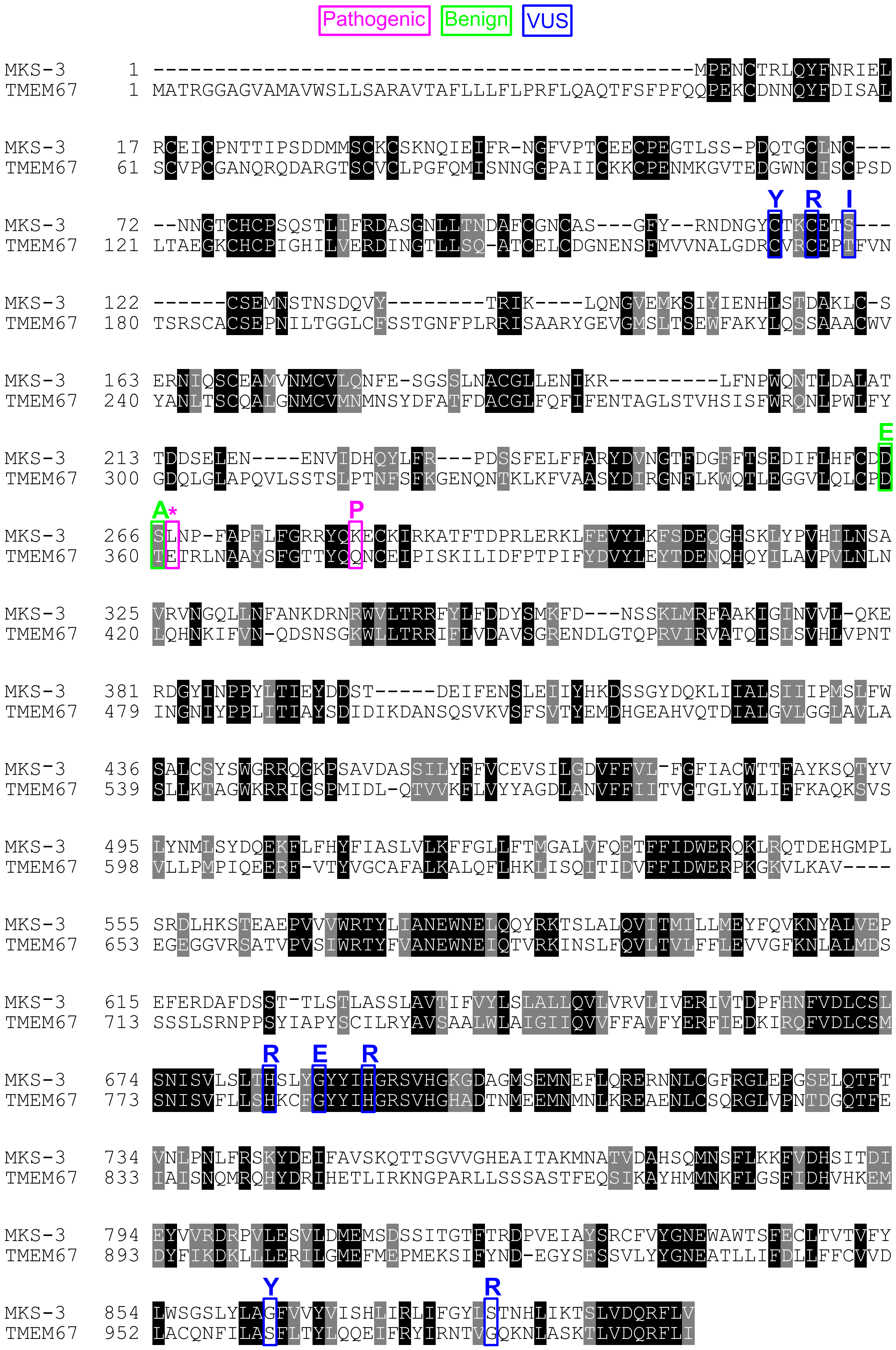

**Figure S1. TMEM67 is conserved from humans to worms.**
Protein alignment between human TMEM67 (NP_714915) and *C. elegans* MKS-3 (NP_495591.2). Conservation of identical (black) and similar (grey) amino acids are highlighted. Amino acids that were mutated in this study are labelled. Amino acid sequences were aligned using Clustal Omega and the figure was generated using BoxShade 3.21.

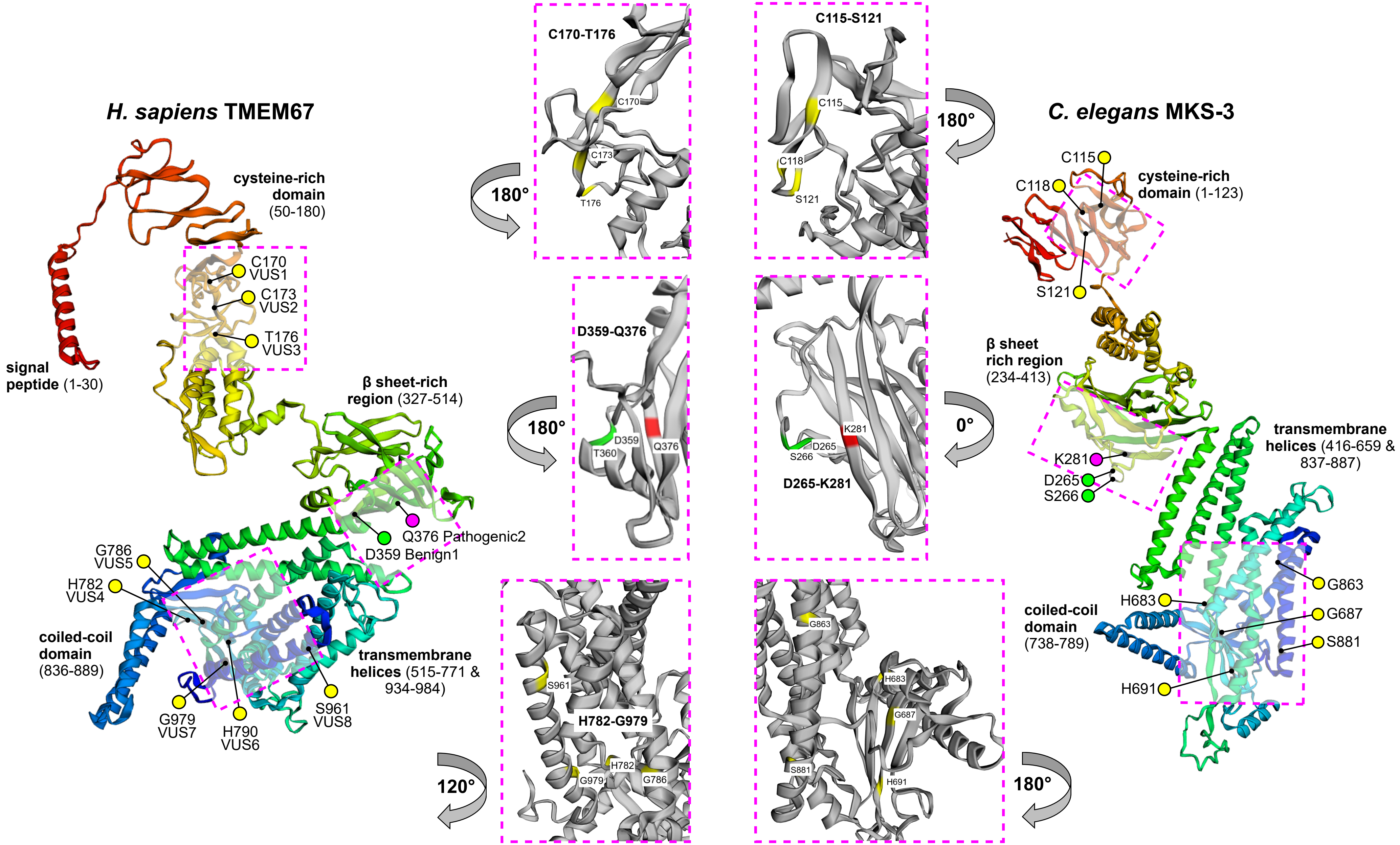

**Figure S2. RaptorX predicted structures of TMEM67 and MKS-3**

Ribbon diagrams of proteins are rainbow-coloured (red at N-terminus to dark blue at C-terminus) with variants indicated (red, known pathogenic; yellow, VUS; green, known benign). Insets highlight amino acids analyzed in this study.

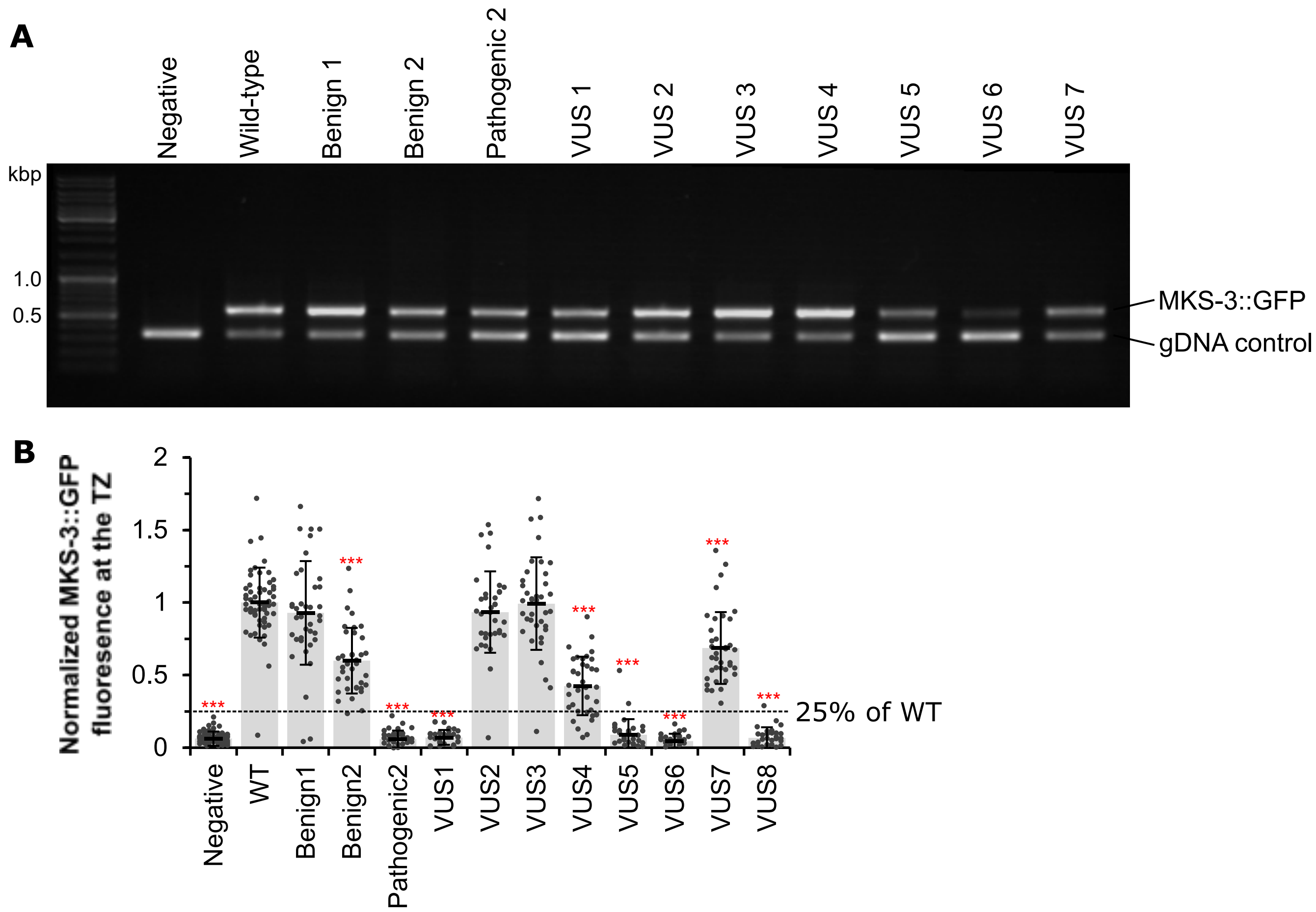

**Figure S3. Transgenic MKS-3::GFP**

A) PCR products after DNA gel electrophoresis. The upper band is specific to *mks-3::gfp*. The lower band is a gDNA control. Non-injected worms (Negative) do not contain the *mks-3::gfp* product while the transgenic strains do. PCR for *VUS8::gfp* is not shown.

B) Quantification of MKS-3::GFP levels at the transition zone. Background fluorescence was subtracted. Individual dots show each measurement while the bars show the average +/- the standard deviations. If a measurements had more than 25% of the wild-type level (dashed line) we concluded that it was positive for MKS-3::GFP localization to the transition zone. Statistical significance according to a one way ANOVA followed by Tukey's *post hoc* test and is relative to the wild-type control.

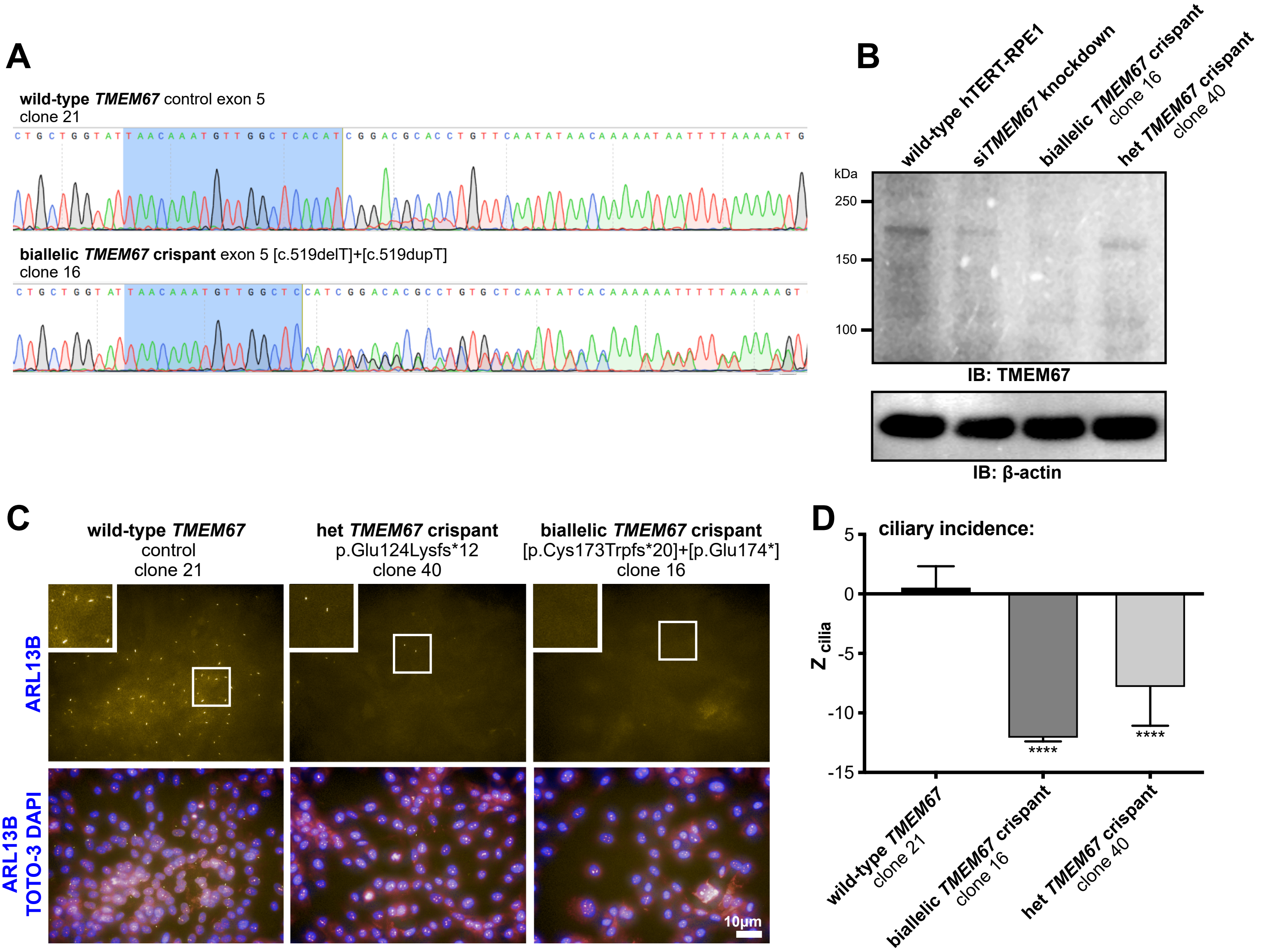

**Figure S4. Characterization of *TMEM67* crispant**

**A)** Sanger sequencing electropherograms comparing *TMEM67* exon 5 sequence between wild-type negative control cell-line clone 21 and the bi-allelic crispant cell-line clone 16. Highlighted sequence in blue indicates the guide RNA sequence used for targeting exon 5. Sequence analysis reveals a one base-pair insertion on one strand, and a one base-pair insertion at the same position on the other strand, corresponding to biallelic frameshift variants: c.519delT, p.(Cys173Trpfs*20) and c.519dupT, p.(Glu174*).

**B)** Western blotting of protein lysates from untreated wild-type hTERT-RPE-1, wild-type hTERT-RPE-1 following siRNA knockdown of *TMEM67*, the bi-allelic *TMEM67* crispant clone 16 and the heterozygous *TMEM67* crispant clone 40, with b-actin as a loading control. A band visible at ca. 200kDa likely corresponds to post-translationally modified TMEM67 (expected molecular weight 112kDa). Levels of TMEM67 expression are most reduced in the bi-allelic knockout crispant clone 16 (no band visible), with decreased levels observed in the heterozygous crispant clone 40 and the siRNA TMEM67 knockdown compared to wild-type.

**C)** hTERT-RPE-1 cells imaged using an "Operetta" (Perkin-Elmer) high-content imaging system, with representative images from Harmony/Columbus software cilia recognition protocol “find spots”. Images show cells stained for the ciliary membrane protein ARL13B (gold), nuclei with DAPI (blue) and cytoplasm with TOTO3 (pink). Significantly fewer cilia were present in the heterozygous *TMEM67* crispant clone 40 compared to wild-type hTERT-RPE-1, and no cilia are visible in the bi-allelic *TMEM67* crispant clone 16. Frames indicate the position of magnified insets. Scale bar = 10mM.

**D)** Bar graphs showing mean robust z score for % ciliated cells (zcilia) for wild-type hTERT-RPE-1 (+0.60), the bi-allelic *TMEM67* crispant clone 16 (-12.15) and the heterozygous *TMEM67* crispant clone 40 (-7.89). Ciliary incidence is significantly decreased (zcilia < -2.0) in both crispant clones compared to wild-type.

**Figure S5. Prediction of deleteriousness of missense alleles using *in silico* analysis**
For each nonsynonymous variant, the text colour denotes if the prediction is tolerated/benign (green), deleterious/damaging (red), or possibly damaging (black). Using these prediction tools we ranked the variants for their overall predicted deleteriousness (1 = benign, 11 = most severe/pathogenic). Variants that are predicted to be more deleterious/damaging are assumed to correlate with disease pathogenesis. The different analyses consistently revealed Benign1 and Benign2 (green background) as benign and the least damaging of all 11 missense mutations. Pathogenic2 (gray background) is identified as deleterious by four of the five prediction tools and ranks as the 3rd most deleterious variant. Overall the *in silico* predictions suggest all eight VUS alleles are likely deleterious/damaging. The only exceptions are VUS3/8 (predicted as probably not damaging by PolyPhen-2 and CADD) and VUS7 (predicted to be tolerated by SIFT).

| TMEM67[a] | Clinical Significance | MISTIC[b] | SIFT[c] | Poly-Phen2[d] | CADD[e] | REVEL[f] | Overall Rank |
|---|---|---|---|---|---|---|---|
| D359E | Benign1 | 0.293 | 0.27 | 0.25 | 17.0 | 0.338 | 1 |
| T360A | Benign2 | 0.437 | 0.18 | 0.55 | 23.2 | 0.476 | 2 |
| Q376P | Pathogenic2 | 0.965 | 0.14 | 0.998 | 26.0 | 0.935 | 9 |
| C170Y | VUS1 | 0.926 | 0 | 1 | 27.1 | 0.889 | 6 |
| C173R | VUS2 | 0.955 | 0 | 1 | 27.2 | 0.926 | 10 |
| T176I | VUS3 | 0.782 | 0.03 | 0.681 | 24.2 | 0.596 | 4 |
| H782R | VUS4 | 0.735 | 0 | 0.959 | 25.3 | 0.927 | 5 |
| G786E | VUS5 | 0.746 | 0 | 1 | 32.0 | 0.919 | 8 |
| H790R | VUS6 | 0.962 | 0 | 1 | 25.7 | 0.988 | 11 |
| G979R | VUS7 | 0.901 | 0.25 | 1 | 28.8 | 0.958 | 7 |
| S961Y | VUS8 | 0.741 | 0.02 | 0.495 | 24.8 | 0.781 | 3 |

a. Amino acid residues correspond to TMEM67 reference sequence NP_714915.
b. MISTIC(MISsense deleTeriousness predICtor) values are from 0-1 with >0.5 being deleterious (Chennen et al. 2020).
c.SIFT(Sorting Intolerant from Tolerant) scores probability of deleteriousness with <0.05 considered significant (Sim et al. 2012). Scores were calculated using an alignment of human TMEM67 with orthologs from mouse, rat, zebrafish, and nematodes.
d. PolyPhen-2(Polymorphism Phenotyping v2) scores range from 0-1 with >0.908 being probably damaging, > 0.446 ≤ 0.908 possibly damaging, and ≤ 0.446 benign (Adzhubei et al. 2010).
e. CADD(Combined Annotation Dependent Depletion) v1.6 phred scores range from 1-99 with higher scores being more deleterious (Rentzsch et al. 2019). We used a cut off of 25.0 because this was the score between the known benign and pathogenic scores.
f. REVEL(Rare Exome Variant Ensemble Learner) scores range from 0-1 with >0.5 being likely pathogenic (Ioannidis et al. 2016).

**Table S1. Variants analyzed in this study**

| | Variant | Clinical Significance | GRCh37 | cDNA | Protein | Additional information | Conditions | Publications |
|---|---|---|---|---|---|---|---|---|
| Benign1 | D359E | --- | Chr8:94794634 | NM_153704.6:c.1077C>G | p.(Asp359Glu) | | | |
| Benign2 | T360A | Likely Benign | Chr8:94794635 | NM_153704.6:c.1078A>G | p.(Thr360Ala) | | Joubert syndrome | Consugar, et al. (2015) Genetics in Medicine 17 (4): 253–61. |
| Pathogenic1 | E361* | Pathogenic | Chr8:94794638 | NM_153704.6:c.1081G>T | p.(Glu361Ter) | in trans with 5 nt insertion/frameshift | Joubert syndrome | Fleming, et al. (2017) CJASN 12 (12), 1962–1973. |
| Pathogenic2 | Q376P | Pathogenic | Chr8:94794684 | NM_153704.6:c.1126C>G | p.(Gln376Pro) | Homozygous | Joubert syndrome | Smith et al. (2006) Nature Genetics 38 (2): 191–96. |
| VUS1 | C170Y | VUS* | Chr8:94777684 | NM_153704.5:c.509G>A | p.(Cys170Tyr) | identified in trans with an established pathogenic splice variant (c.224-2A>T) | Meckel syndrome | This study. |
| VUS2 | C173R | VUS | Chr8:94777644 | NM_153704.6:c.517T>C | p.(Cys173Arg) | Compound heterozygote with S312P | Meckel syndrome | Huynh, et al. (2018) Clinical Case Reports 6(11), 2189–2192. |
| VUS3 | T176I | VUS* | Chr8:94777654 | NM_153704.5:c.527C>T | p.(Thr176Ile) | identified *in trans* with an established pathogenic variant | Meckel syndrome | This study. |
| VUS4 | H782R | VUS | Chr8:94817012 | NM_153704.4:c.2345A>G | p.(His782Arg) | | COACH | Brancati, et al. (2009) Human Mutation 30 (2): E432–42. |
| VUS5 | G786E | VUS | Chr8:94817024 | NM_153704.6:c.2357G>A | p.(Gly786Glu) | Biallelic with nonsense allele (E848*) | Meckel syndrome | Iannicelliet al. (2010) Human Mutation 31(5), E1319–E1331. |
| VUS6 | H790R | VUS* | Chr8:94817036 | NM_153704.5:c.2369A>G | p.(His790Arg) | Identified heterozygous in a parent | Meckel syndrome | This study. |
| VUS7 | G979R | VUS* | Chr8:94828627 | NM_153704.5:c.2935G>A | p.(Gly979Arg) | identified in trans with an established pathogenic splice variant(c.507-3C>A) | Meckel syndrome | This study. |
| VUS8 | S961Y | VUS/Likely pathogenic | Chr8:94827650 | NM_153704.5:c.2882C>A | p.(Ser961Tyr) | identified *in trans* with an established pathogenic variant, no functional evidence | Meckel Syndrome | Szymanska, K., et al. (2012). Cilia, 1 (18). |

*VUS1(C170Y), VUS3(T176I), VUS6(H790R), and VUS7(G979R) were identified from whole exome sequencing of suspected Meckel syndrome feotuses.

**Table S2. Worm strains**

| | Strain | Genotype | Details |
|---|---|---|---|
| Controls | N2 | Wild-type | |
| | | *nphp-4(tm925) V* | |
| | | *mks-3(tm2547) II* | 949 bp deletion |
| | | *mks-3(tm2547) II; nphp-4(tm925) V* | |
| | | *dpy-5(e907) I; mks-3(tm2547) II; nphp-4(tm925) V* | Used to generate heterozygotes |
| CRISPR mutants | OEB934 | *mks-3(oq123[K281P]) II; nphp-4(tm925) V* | Pathogenic 2, K281P |
| | OEB935 | *mks-3(oq124[D265E]) II; nphp-4(tm925) V* | Benign 1, D265E |
| | OEB941 | *mks-3(oq130[L267*]) II; nphp-4(tm925) V* | Pathogenic 1, L267* |
| | OEB942 | *mks-3(oq131[S266A]) II; nphp-4(tm925) V* | Benign 2, S266A |
| | OEB943 | *mks-3(oq132[C115Y]) II; nphp-4(tm925) V* | VUS1, C115Y |
| | OEB944 | *mks-3(oq133[C118R]) II; nphp-4(tm925) V* | VUS2, C118R |
| | OEB955 | *mks-3(oq134[H683R]) II; nphp-4(tm925) V* | VUS4, H683R |
| | OEB956 | *mks-3(oq135[G687E]) II; nphp-4(tm925) V* | VUS5, G687E |
| | OEB957 | *mks-3(oq136[H691R]) II; nphp-4(tm925) V; unc-58(oq146) X* | VUS6, H691R |
| | OEB1018 | *mks-3(oq136[H691R]) II; nphp-4(tm925) V* | VUS6, H691R, outcrossed 1x |
| | OEB980 | *mks-3(oq139[S121I]) II; nphp-4(tm925) V* | VUS3, S121I |
| | OEB981 | *mks-3(oq140[S881R]) II; nphp-4(tm925) V; unc-58(oq147) X* | VUS7, S881R |
| | OEB1019 | *mks-3(oq140[S881R]) II; nphp-4(tm925) V* | VUS7, S881R, outcrossed 1x |
| | OEB1001 | *mks-3(oq145[G863Y]) II; nphp-4(tm925) V* | VUS8, G863Y |
| Extrachromosomal Arrays | OEB990 | *mks-3(tm2547) II; oqEx122[mks-3p::mks-3::gfp + coel::dsRed]* | Wild-type |
| | OEB991 | *mks-3(tm2547) II; oqEx123[mks-3p::mks-3(oq124)::gfp + coel::dsRed]* | Benign 1, D265E |
| | OEB992 | *mks-3(tm2547) II; oqEx124[mks-3p::mks-3(oq131)::gfp + coel::dsRed]* | Benign 2, S266A |
| | OEB993 | *mks-3(tm2547) II; oqEx125[mks-3p::mks-3(oq123)::gfp + coel::dsRed]* | Pathogenic 2, K281P |
| | OEB994 | *mks-3(tm2547) II; oqEx126[mks-3p::mks-3(oq132)::gfp + coel::dsRed]* | VUS1, C115Y |
| | OEB995 | *mks-3(tm2547) II; oqEx127[mks-3p::mks-3(oq133)::gfp + coel::dsRed]* | VUS2, C118R |
| | OEB996 | *mks-3(tm2547) II; oqEx128[mks-3p::mks-3(oq139)::gfp + coel::dsRed]* | VUS3, S121I |
| | OEB997 | *mks-3(tm2547) II; oqEx129[mks-3p::mks-3(oq134)::gfp + coel::dsRed]* | VUS4, H683R |
| | OEB998 | *mks-3(tm2547) II; oqEx130[mks-3p::mks-3(oq135)::gfp + coel::dsRed]* | VUS5, G687E |
| | OEB999 | *mks-3(tm2547) II; oqEx131[mks-3p::mks-3(oq136)::gfp + coel::dsRed]* | VUS6, H691R |
| | OEB1000 | *mks-3(tm2547) II; oqEx132[mks-3p::mks-3(oq140)::gfp + coel::dsRed]* | VUS7, S881R |
| | OEB1020 | *mks-3(tm2547) II; oqEx133[mks-3p::mks-3(oq145)::gfp + coel::dsRed]* | VUS8, G863Y |

### Table S3. Worm crRNA sequences and repair templates

| crRNA sequence | Allele | Mutation | ssODN Repair Template Sequence* |
|---|---|---|---|
| ATCCACGCACATGGTCACTA | --- | *unc-58* co-CRISPR | atttgtggtataaaatagccgagttaggaaacaaatttttctttcagGTtTTTCtGTcGTtACCATGTGCGTGGATCTTGCGTCCACACATCTCAAGGCGTACTT |
| AAAAGAATGCAAAATAAGAA | *oq123* | Pathogenic 2, K281P | TGAACCCATTTGCTCCATTTTTATTCGGAAGACGTTATCAAccAGAgTGCAAgATccGtAAGGCAACATTCACAGATCCACGTCTTGAGAGAAAATTATT |
| CGAATAAAAATGGAGCAAAT | *oq124* | Benign 1, D265E | TTTTTCACTTCGGAAGACATATTTCTTCATTTTTGTGATGAgTCcCTtAAtCCATTcGCcCCATTTTTATTCGGAAGACGTTATCAAAAAGAATGCAAAAT |
| | *oq130* | Pathogenic 1, L267* | TTTCACTTCGGAAGACATATTTCTTCATTTTTGTGATGACTCTtgaAAtCCATTcGCcCCATTTTTATTCGGAAGACGTTATCAAAAAGAATGCAAAATAA |
| | *oq131* | Benign 2, S266A | TTTTCACTTCGGAAGACATATTTCTTCATTTTTGTGATGACgCcCTcAAtCCATTcGCcCCATTTTTATTCGGAAGACGTTATCAAAAAGAATGCAAAAT |
| GGCTTTTACAGAAATGACAA | *oq132* | VUS1, C115Y | gaatgaacaaacCATTTCGGAACACGACGTTTCACATTTCGTatAgTATCCGTTaTCgTTTCTGTAAAAGCCAGAAGCACAGTTTCCACAGAATGCATCA |
| | *oq133* | VUS2, C118R | atgtgaatgaacaaacCATTTCGGAACACGACGTTTCACgcTTgGTGCAgTATCCGTTaTCgTTTCTGTAAAAGCCAGAAGCACAGTTTCCACAGAATGCA |
| TCTCTGACCCATTCTCTGTA | *oq134* | VUS4, H683R | CATCACCCTTTCCATGAACAGAACGACCATGAATATAGTATCCATAgAGgGAgcGGGTCAGAGAAAGGACActgaaattgatttaagccgtgaagaattt |
| | *oq135* | VUS5, G687E | CCAGCATCACCCTTTCCATGAACAGAACGACCATGAATATAGTActCATAgAGgGAgTGGGTCAGAGAAAGGACActgaaattgatttaagccgtgaagaa |
| | *oq136* | VUS6, H691R | TCATTCCAGCATCACCCTTTCCATGAACAGAACGACCAcGgATgTAGTAaCCATAgAGgGAgTGGGTCAGAGAAAGGACActgaaattgatttaagccgt |
| TGTGAATGAACAAACCATTT | *oq139* | VUS3, S121I | GCTTTTACAGAAATGACAACGGATATTGCACGAAATGTGAAACGatcTGcTCtGAgATGgtttgttcattcacattttagtgttttctttttggaatact |
| ACAAGTAACCGAAAATCAAA | *oq140* | VUS7, S881R | TGCTGGTTTTGTTGTGTATGTTATTTCTCACTTGATCCGccTcATcTTCGGaTACcTccgtACGAATCATTTGATTAAGACGAGTTTAGTAGATCAACGA |
| TCTGGATCTTTATATCTTGC | *oq145* | VUS8, G863Y | ATGTTTAACAGTTACAGTCTTCTATTTATGGTCTGGATCTTTATAcCTcGCTtacTTTGTTGTGTATGTTATTTCTCACTTGATCCGTTTGATTTTCGGT |

*lowercase red nucleotides are engineered mutations

### Table S4. Worm sequencing/PCR primers

| ID | Name | Sequence | Purpose |
|---|---|---|---|
| NL132 | F35D2.4_R+2720 | tggttcaagtcctcggaatc | Genotype *tm2547* |
| NL446 | F35D2.4_F+374 | tgcccttcacaaagcactct | Genotype *tm2547* |
| NL447 | F35D2.4_F+1537 | catcatattcaattgttaaatacggtg | Genotype *tm2547* |
| KL154 | mks-3.F-2 | ctATGCCTGAAAATTGTACGAG | Genotype *oq132, oq133* |
| KL155 | mks-3.R+718 | tcaaaatgaaacCTCGCAAC | Genotype *oq132, oq133, oq139* |
| KL156 | mks-3.C115Y | CGACGTTTCACATTTCGTatAg | Genotype *oq132* |
| KL159 | mks-3.C118R | AACACGACGTTTCACgcTTg | Genotype *oq133* |
| KL161 | mks-3.F+798 | TTCACTGAATGCGTGTGGAC | Genotype *oq123, oq124, oq130, oq131* |
| KL162 | mks-3.R+1664 | AGAGCTGACCAGAACAATGAC | Genotype *oq123, oq131* |
| KL163 | mks-3.D265E | gGCgAATGGaTTaAGgGAc | Genotype *oq124* |
| KL164 | mks-3.S266A | CATTTTTGTGATGACgCcCTc | Genotype *oq131* |
| KL165 | mks-3.L267Ter | TAAAAATGGgGCgAATGGaTTtc | Genotype *oq130* |
| KL166 | mks-3.K281P | CTTaCggATcTTGCAcTCTgg | Genotype *oq123* |
| KL176 | mks-3.R.1123 | AAATGGAGCAAATGGGTTC | Genotype *oq124, oq130, oq131* |
| KL177 | mks-3.For-39 | CAAATGCTCAGTTTCGTTCAC | Genotype *oq139* |
| KL178 | mks-3.F+2936 | TCGGGATCGACCAGTCTTG | Genotype *oq140* |
| KL179 | mks-3.R+3649 | caggagatcagtgcgaacg | Genotype *oq140* |
| KL181 | mks-3.F+2125 | TGCGAATGAATGGAATGAAC | Genotype *oq134, oq135* |
| KL182 | mks-3.R+2814 | CAGATGTAGTTTGTTTGGAGAC | Genotype *oq134, oq135, oq136* |
| KL184 | mks-3.S881R-Rev | TAATCAAATGATTCGTacggAg | Genotype *oq140* |
| KL185 | mks-3.H683R-Rev | ATAGTATCCATAgAGgGAgc | Genotype *oq134* |
| KL186 | mks-3.G687E-For | GACCCAcTCcCTcTATGag | Genotype *oq135* |
| KL188 | mks-3.H691R-For | cTATGGtTACTAcATcCg | Genotype *oq136* |
| KL199 | mks-3.For+2532 | TTCTCTGACCCATTCTCTG | Genotype *oq136* |
| KL200 | S121I.Rev | caaacCATcTCaGAgCAgat | Genotype *oq139* |
| KL183 | mks-3.G863Y-Rev | TACACAACAAAgtaAGCgAGg | Genotype *oq145* |
| KL230 | unc-58. For+3669 | GACTCGGAGATATCGTTGTGACTG | *unc-58* PCR |
| KL231 | unc-58. Rev+4393 | CGCGGAGTTCGTTATCCAGGAAG | *unc-58* PCR |
| KL232 | unc-58. Rev+4367 | CGCACATCATTCCATGTAAC | Sequencing primer |
| KL229 | mks-3.For-495 | tgtctttgactaggcataacccaac | *mks-3::gfp* stitch (PCR1) |
| NL441 | F35D2.4_F-1196 | tggtaatttgctcagtgtttcaattg | *mks-3::gfp* stitch (PCR1) |
| NL443 | mks-3_GFP_R1 | GAGTCGACCTGCAGGCATGCAAGCTTaacaagaaatcgttgatctactaaactcgt | *mks-3::gfp* stitch (PCR1) |
| ST54 | mks-3_-485_F | taggcataacccaacaatcaac | *mks-3::gfp* stitch (PCR2) |
| NL74 | GFP Rev (D*) | GGAAACAGTTATGTTTGGTATATTGGG | *mks-3::gfp* stitch (PCR2) |

**Table S5. Site Directed Mutagenesis Primers**

| Target TMEM67 cDNA change | Target TMEM67 protein change | Primer Direction | Sequence (5' to 3') |
|---|---|---|---|
| c.509G>A | Cys170Tyr | Forward | gctttaggagacaggtacgtccgatgtgagc |
| | | Complement | gctcacatcggacgtacctgtctcctaaagc |
| c.515G>A | Arg172Gln | Forward | aaatgttggctcacattggacgcacctgtctcc |
| | | Complement | ggagacaggtgcgtccaatgtgagccaacattt |
| c.517T>C | Cys173Arg | Forward | caaatgttggctcacgtcggacgcacctgtc |
| | | Complement | gacaggtgcgtccgacgtgagccaacatttg |
| c.527C>T | Thr176Ile | Forward | ctgctggtattaacaaatattggctcacatcggacgc |
| | | Complement | gcgtccgatgtgagccaatatttgttaataccagcag |
| c.1077C>G | Asp359Glu | Forward | ccttgtctctgtctctggacaaagctgtaaaacac |
| | | Complement | gtgttttacagctttgtccagagacagagacaagg |
| c.1078A>G | Thr360Ala | Forward | catttagccttgtctctgcgtctggacaaagctgtaa- |
| | | Complement | ttacagctttgtccagacgcagagacaaggctaaatg |
| c.2935G>A | Gly979Arg | Forward | atgccaaattcttttgtcttactgtattacggatatatctaaaaatctcttgt |
| | | Complement | acaagagatttttagatatatccgtaatacagtaagacaaaagaatttggcat |
| c.2882C>A | Ser961Tyr | Forward | tcttgttgtagatatgtaaggaagtatgctaaaataaaattttggcaagc |
| | | Complement | gcttgccaaaattttattttagcatacttccttacatatctacaacaaga |
| c.1127A>C | Gln376Pro | Forward | gagataggaatctcacaatttggttggtaggttgttccaaatg |
| | | Complement | catttggaacaacctaccaaccaaattgtgagattcctatctc |

Primers were designed using the web-based QuikChange Primer Design Program (https://www.agilent.com/store/primerDesignProgram.jsp).

### Table S6. TMEM67 Alt-R crRNAs

| TMEM67 target exon | crRNA sequence | PAM | Specificity Score | Efficiency Score |
|---|---|---|---|---|
| 2 | CAGATGATCTCTAATAATGG | AGG | 63.16 | 71.18 |
| 3 | CCTAGTGACTTAACTGCCGA | AGG | 90.6 | 63.58 |
| 5 | TAACAAATGTTGGCTCACAT | CGG | 64.11 | 71.28 |

| Table S7. TMEM67 sequencing/PCR primers | | |
|---|---|---|
| **Target** | **Primer name** | **Primer sequence (5' to 3')** |
| Internal TMEM67 | cDNA1-RV | AACAGTGCTCAGTCCAGCAG |
| Internal TMEM67 | EXON8R | CACACATATTTCCAAGAGCTTGAC |
| Internal TMEM67 | EX4-7R | TTGCAAACCATTCTGAAGTTAAAG |
| Internal TMEM67 | EX4-7F | TTGTGAGCTCTGTGATGGAAA |
| Internal TMEM67 | EXON3F | TGTCCCATTGGCCATATTTT |
| Internal TMEM67 | EX27-3’UTR-R | ACTACACACAATGGGAAAACAGTA |
| Internal TMEM67 | ISH2R | AAAAATATGGCAAACTAACCTGA |
| Internal TMEM67 | CDNA5-RV | AAAAATATGGCAAACTAACCTGA |
| Internal TMEM67 | EXONIC 2F | TGTAAAAAGTGCCCAGAAAACA |
| Internal TMEM67 | EX27-3’UTR-F | TGTGTTGTGGATTTGGCTTG |
| Internal TMEM67 | ISH3F | TCTTGGCTCCTTCATTGACC |
| Internal TMEM67 | CDNA4-RV | TGGGTACCAAACCTCTCTGG |
| Internal TMEM67 | ISH2F | TCGACAGTTCGTTGATTTATGC |
| Internal TMEM67 | EXON20/21R | CCCACAACCTCCAAAAAGAA |
| Internal TMEM67 | EX19R | GAGCTTATGCAAAAATTGTAGTGC |
| Internal TMEM67 | ISH1R | AGACTGGCTGTTGGCATCTT |
| Internal TMEM67 | CDNA2-RV | TCGAATTACTCTTGGCTGAGTTC |
| Internal TMEM67 | ISH6F | TTTTGGCTGTGCCTGTGTTA |
| Internal TMEM67 | ISH5R | GGAAGCAGCAACAAACTTCA |
| Internal TMEM67 | TMEM67_1203_F | GGCTGTGCCTGTGTTAAACC |
| Internal TMEM67 | TMEM67_1481_R | GACTGGCTGTTGGCATCTTTG |
| Internal TMEM67 | TMEM67_1942_F | GTACGAAGTGCCACTGTTCCTG |
| Internal TMEM67 | TMEM67_2459_R | GTCTGACCATCTGTGTTGGGT |
| Internal PX458 | T7 | TAATACGACTCACTATAGGG |
| Internal PX458 | U6 | GACTATCATATGCTTACCGT |
| Internal PX458 | BGHR | TAGAAGGCACAGTCGAGG |
| Internal PX458 | CMV-Forward | CGCAAATGGGCGGTAGGCGTG |
| Internal PX458 | EGFP-C-For | CATGGTCCTGCTGGAGTTCGTG |
| Internal PX458 | EGFP-C-REV | GTTCAGGGGGAGGTGTG |
| Internal PX458 | EGFP-N | CGTCGCCGTCCAGCTCGACCA |
| Internal PX458 | EXFP-R | GTCTTGTAGTTGCCGTCGTC |
| Internal PX458 | F1ori-F | GTGGACTCTTGTTCCAAACTGG |
| Internal PX458 | M13 F | TGTAAAACGACGGCCAGT |
| Internal PX458 | pBR322ori-F | GGGAAACGCCTGGTATCTTT |
| Internal PX458 | SP6 | ATTTAGGTGACACTATAG |
| Internal PX458 | T7 | TAATACGACTCACTATAGGG |
| Internal PX458 | SpCas9_1F | GCCAAGGTGGACGACAGCTT |
| Internal PX458 | SpCas9_2F | ACCTACAACCAGCTGTTCGAGG |
| Internal PX458 | SpCas9_3F | CAAGAACCTGTCCGACGCC |
| Internal PX458 | SpCas9_4F | GTGAAGCTGAACAGAGAGGACCT |
| Internal PX458 | SpCas9_5F | TTCATCGAGCGGATGACCAACT |
| Internal PX458 | SpCas9_6F | CTGGGCACATACCACGATCTG |
| Internal PX458 | SpCas9_7F | AACTTCATGCAGCTGATCCACGA |

| | | |
|---|---|---|
| Internal PX458 | SpCas9_8F | GGCAGCCAGATCCTGAAAGAAC |
| Internal PX458 | SpCas9_9F | CGCCAAGCTGATTACCCAGAG |
| Internal PX458 | SpCas9_10F | GTGGGAACCGCCCTGATCAA |
| Internal PX458 | SpCas9_11F | TCGTGAAAAAGACCGAGGTGC |
| Internal PX458 | SpCas9_12F | AGGACCTGATCATCAAGCTGC |
| Internal PX458 | SpCas9_13F | GTCCGCCTACAACAAGCACC |
| Internal PX458 | Cas9_14_F | GCTTCACTCTCCCCATCTCC |
| Internal PX458 | Cas9_15_F | GCGCTCCGAAAGTTTCCTTT |
| TMEM67 CRISPR targets | Exon2_F | CCCTAGTCCCCGTAAATGGA |
| TMEM67 CRISPR targets | Exon3_F | CTGGTTCAAGCCATTCTGGT |
| TMEM67 CRISPR targets | Exon5_F | CCCTCCCTCTGTCTCATTCA |
| TMEM67 CRISPR targets | Exon 2_R | ACACATCCCAGACTGCTCTA |
| TMEM67 CRISPR targets | Exon3_R | GGCTGAGCAACATAACGAGA |
| TMEM67 CRISPR targets | Exon5_R | AACAGGCTACGGTACAGACT |
